# Availability of abundant thiamine determines efficiency of thermogenic activation in human neck area derived adipocytes

**DOI:** 10.1101/2022.05.05.490662

**Authors:** Rini Arianti, Boglárka Ágnes Vinnai, Ferenc Győry, Andrea Guba, Éva Csősz, Endre Kristóf, László Fésüs

**Author notes:** Equally contributed and corresponding authors. Electronic address.

## Abstract

Brown/beige adipocytes express uncoupling protein-1 (UCP1) that enables them to dissipate energy as heat. Systematic activation of this process can alleviate obesity. Human brown adipose tissues are interspersed in distinct anatomical regions including deep neck. We found that UCP1 enriched adipocytes differentiated from precursors of this depot highly expressed ThTr2 transporter of thiamine and consumed thiamine during thermogenic activation of these adipocytes by cAMP which mimics adrenergic stimulation. Inhibition of ThTr2 led to lower thiamine consumption with decreased proton leak respiration reflecting reduced uncoupling. In the absence of thiamine, cAMP-induced uncoupling was diminished but restored by thiamine addition reaching the highest levels at thiamine concentrations larger than present in human blood plasma. Thiamine is converted to thiamine pyrophosphate (TPP) in cells; the addition of TPP to permeabilized adipocytes increased uncoupling fueled by TPP-dependent pyruvate dehydrogenase. ThTr2 inhibition also hampered cAMP-dependent induction of UCP1, PGC1a, and other browning marker genes, and thermogenic induction of these genes was potentiated by thiamine in a concentration dependent manner. Our study reveals the importance of amply supplied thiamine during thermogenic activation in human adipocytes which provides TPP for TPP-dependent enzymes not fully saturated with this cofactor and by potentiating the induction of thermogenic genes.

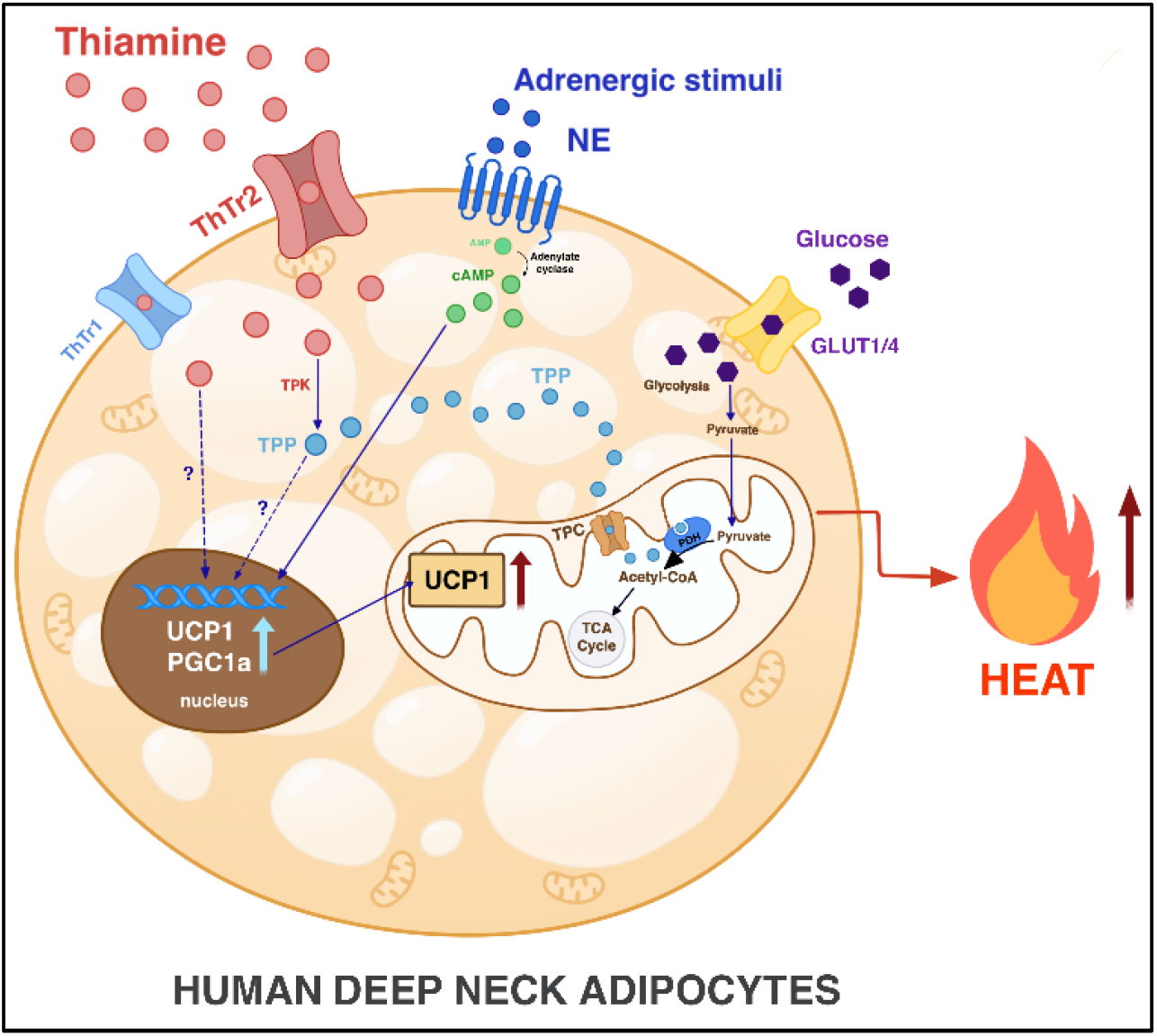

**Highlights:**

- Abundant thiamine is required for efficient activation of UCP1 dependent thermogenesis in human deep neck derived adipocytes
- Inhibition of thiamine transporters leads to decreased thermogenic response
- In stimulated adipocytes, thiamine supply provides extra thiamine pyrophosphate for increasing pyruvate dehydrogenase activity to generate sufficient fuel of UCP1 dependent respiration
- Adrenergic stimulation of thermogenic gene expression is potentiated by thiamine in a concentration dependent manner

## Introduction

The highly adoptive and heterogeneous adipose tissue depots have multiple functions which include storage of energy reservoirs, production of hormones that control physiological processes, regulation of inflammatory responses, insulation of body parts, and participation in heat production (Sakers et al., 2022). Thermogenic brown adipose tissues (BAT) mediate energy expenditure via uncoupling protein-1 (UCP1) dependent uncoupling of proton gradient from ATP synthesis to maintain constant core body temperature in newborns and hibernating animals (Townsend and Tseng, 2014). Human BAT was primarily regarded as a tissue which was only present in infants and located at anatomical sites that are difficult to be reached. Positron emission tomography (PET) provided evidence that adult humans also have BAT which can be activated by cold exposure and adrenergic stimulation (Cypess et al., 2009; Virtanen et al., 2009). Brown and variably ‘brownable’ (often described as beige) adipose tissue depots could be located in several anatomical regions, including cervical deep neck (DN), supraclavicular, axillary, paraspinal, and mediastinal depots (Leitner et al., 2017).

Molecular signature of human DN adipose tissue and adipocytes has been extensively investigated in the last decade. Cypess et al. reported that classical brown adipocytes originally described in mice are present in the DN depot marked by high expression of UCP1, ZIC1, and LHX8 (Cypess et al., 2013). Others have found that human brown adipocytes derived from DN or supraclavicular area have more resemblance to murine beige adipocytes (Wu et al., 2012; Shinoda et al., 2015) while there are evidences that the molecular signature of classical brown and beige adipocytes are partially overlapping in DN derived adipocytes (Jespersen et al., 2013; Sanchez-Gurmaches et al., 2016). Functional BAT isolated with the help of PET-CT scan guidance from supraclavicular region after cold exposure could be characterized by a large number of genes expressed differentially in comparison to subcutaneous (SC) white adipose tissue (WAT) (Perdikari et al., 2018). Our previous study using stromal vascular fraction (SVF) isolated from neck tissues also demonstrated that DN derived, differentiated adipocytes exerted high browning capacity as compared to SC originated ones (Toth et al., 2020) and characteristic pattern of differentially expressed genes (DEGs) revealed DN associated pathways which were upregulated (thermogenesis, interferon, cytokine, and retinoic acid, with *UCP1* as a prominent network stabilizer) or downregulated (particularly extracellular matrix remodeling) upon brown differentiation. In the presence of the FTO rs1421085 obesity-risk allele, the expression of mitochondrial and thermogenesis genes was suppressed (Toth et al., 2020; Claussnitzer et al., 2015).

The augmentation of heat generation by brown/beige adipocytes has been postulated as one of the promising approaches to treat obesity. They play an important role as metabolic sink for glucose, fatty acids, and branched chain amino acids (Yoneshiro et al., 2019). Enhancing differentiation of progenitor cells towards potentially thermogenic brown/beige adipocytes and stimulating heat generation by the existing ones improve the systemic glucose and lipid homeostasis resulting in a better metabolic health (Harms et al., 2013; Verkerke et al., 2020; Hussain et al., 2020). Abundance of thermogenic adipose tissue in humans is positively correlated with lower body mass index (Cypess et al., 2009). Transplantation of beige differentiated human adipose-derived stromal cells into mice elevated systemic energy expenditure and decreased body mass (Singh et al., 2020). Using a browning gene signature algorithm, it was found that brown adipocyte content in SC WAT increased after caloric restriction and associated with weight loss and energy expenditure (Perdikari et al., 2018). However, designing BAT based anti-obesity protocols in humans requires detailed molecular information about the activation pathways and understanding the long-term maintenance of adipocyte thermogenesis.

Here we report that the expression of 21 solute carrier (SLC) transporters was found differentially regulated in DN versus SC derived adipocytes. Thiamine transporter (ThTr) 2, which is encoded by *SLC19A3*, was among the upregulated SLCs in DN derived adipocytes. Thiamine, the water-soluble vitamin B1, is available in nM concentrations in blood plasma and converted to its active form thiamine pyrophosphate (TPP) by TPP kinase (TPK) in cells (Manzetti et al., 2014). Several metabolic pathways, such as tricarboxylic acid (TCA) cycle and pentose-phosphate cycle require TPP because it acts as a coenzyme for pyruvate dehydrogenase (PDH) (catalyzes the conversion of pyruvate to acetyl-CoA), α-ketoglutarate dehydrogenase (catalyzes the conversion of α-ketoglutarate to succinyl-CoA), transketolase (shifts excess fructose-6-phosphate to glyceraldehyde-3-phosphate) and branched chain α keto-acid dehydrogenase (mediates anaplerosis of TCA). Most of the reactions catalyzed by TPP-dependent enzymes contribute to production of NADH in the mitochondria feeding the electron transport chain to generate the proton gradient. Therefore, thiamine plays a critical role in energy metabolism and thiamine deficiency leads to the accumulation of anaerobic metabolites, such as lactate, and severe neurological symptoms (Manzetti et al., 2014). The prevalence of thiamine deficiency is high in less developed countries of Asia and Africa leading to beriberi, a major cause of death in infants (Whitfield et al., 2018).

Our presented data demonstrate the significance of ThTr2 and the importance of the availability of thiamine during thermogenic activation in human DN derived adipocytes. The inhibition of ThTr2 by its potent inhibitor in the presence of thiamine and thiamine depletion in the cell culture decreased cAMP mediated thermogenic proton leak respiration and halted the cAMP-stimulated induction of thermogenic marker genes in DN derived adipocytes with a less pronounced effect in SC adipocytes. The direct stimulation of the TPP-dependent mitochondrial PDH in permeabilized adipocytes elevated proton leak respiration. The results raise the possibility of using thiamine-dependent regulation mechanism to augment heat generation by thermogenic adipocytes for prevention or treatment of obesity.

## Results

### UCP1 enriched adipocytes express high level of thiamine transporter 2

Previously, we have studied differentiated adipocyte population derived from SVFs of paired DN and SC adipose tissue biopsies of nine donors and compared their global gene expression patterns by RNA sequencing (Toth et al., 2020). The adipocytes were differentiated by regular adipogenic differentiation medium (ADIP) or long-term rosiglitazone treatment (B-ADIP) which enhanced brown differentiation (Toth et al., 2020). There were 1049 DEGs found in DN and SC comparison, and DN derived adipocytes showed higher browning content and capacity quantified by BATLAS and ProFAT scores, respectively (Perdikari et al., 2018; Cheng et al., 2018). Active brown/beige adipocytes consume high amounts of cellular nutrients as well as metabolic cofactors to provide sufficient fuel for heat generation and SLC transporters play a crucial role in mediating their transport into cells. Among the DEGs, we found 21 SLC transporters (Fig. 1a) regulated by either the anatomical location (15 upregulated, 3 downregulated in DN versus SC ADIPs) or the applied differentiation protocol (3 upregulated in both SC and DN B-ADIPs). One of the SLCs genes with higher expression in DN ADIP was thiamine transporter 2, ThTr2 (encoded by *SLC19A3)*, which was undetectable in preadipocytes and was strongly induced during differentiation (Fig. 1b). The expression of ThTr2 was higher in adipocytes derived from DN compared to SC ones, irrespective to the applied differentiation protocol (Fig. 1b) and qPCR analysis with independent donors validated differential expression. Protein levels of ThTr2 detected by immunoblotting were similar to the UCP1 pattern showing that ThTr2 was more expressed in high UCP1 containing DN derived adipocytes (Fig. 1c). ThTr1, encoded by *SLC19A2*, had lower expression level and was not differently expressed between SC and DN derived adipocytes (Fig. 1c, d).

**Figure 1.**
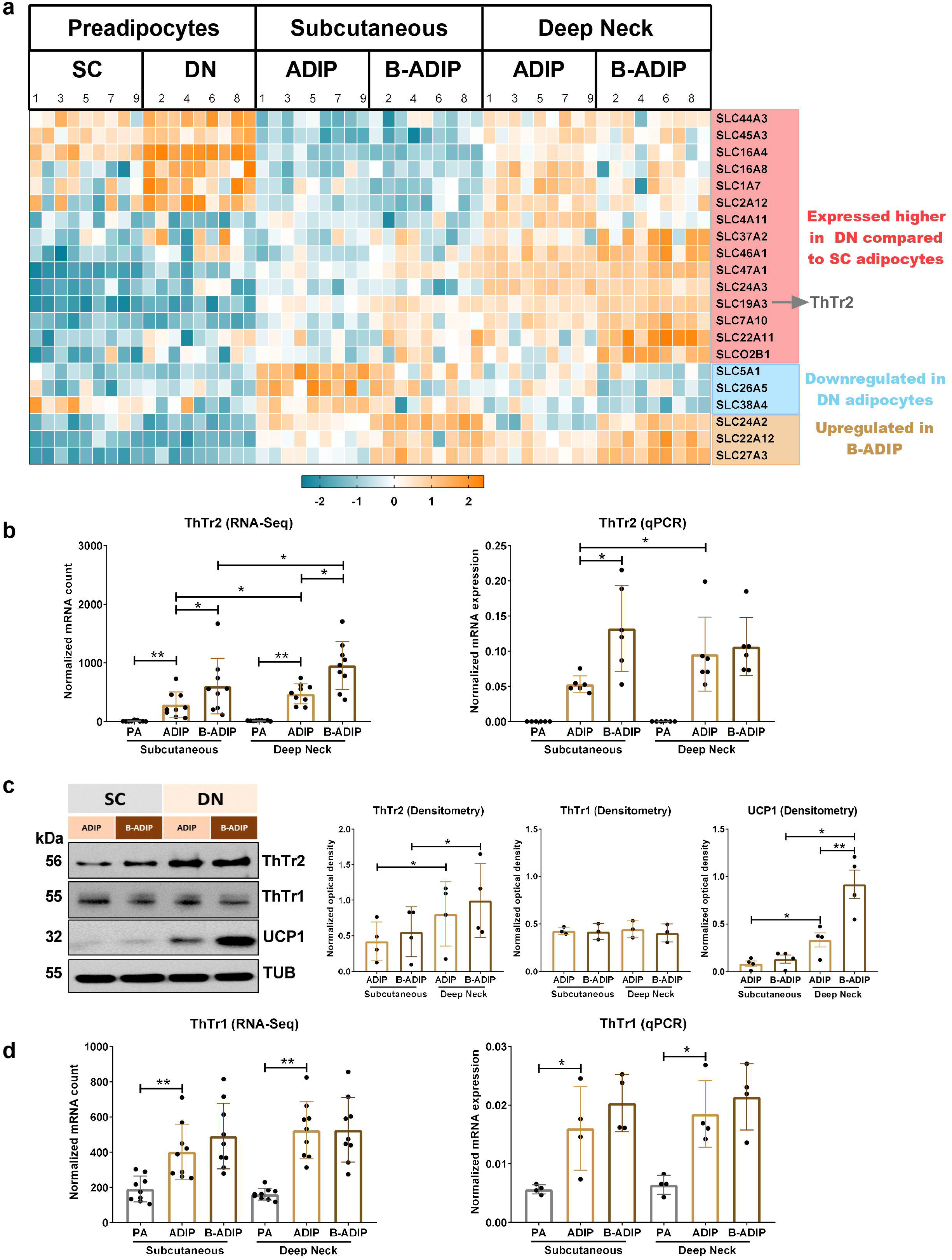
Expression pattern of solute carrier (SLC) and thiamine transporters, ThTr1 and ThTr2 in subcutaneous (SC) and deep neck (DN) adipocytes. SC and DN-derived preadipocytes were differentiated into ADIP and B-ADIP. (a) Heatmap displaying the expression of 21 SLC transporter genes in SC and DN progenitors differentiated to ADIP and B-ADIP. (b) mRNA expression of ThTr2 based on RNA-Seq data, n=9, and assessed by RT-qPCR, n=6. (c) Detection and quantification of ThTr2, ThTr1 and UCP1 protein expression (normalized to tubulin) by immunoblotting, n=4 or 3 (ThTr1) (d) mRNA expression of ThTr1 based on RNA-Seq data, n=9, and assessed by RT-qPCR, n=4. PA: preadipocyte, ADIP: adipocytes. Statistical analysis was performed by unpaired t-test, *p<0.05, **p<0.01 or GLM (b and d RNA seq).

According to a publicly available transcriptomic data set of thermogenic adipocytes derived from progenitors of human abdominal SC WAT (Min et al., 2019), adrenergic stimulation elevated the mRNA expression of both ThTrs (Fig. S1a-b). An RNAseq-based screen identified ThTr2 as adipose tissue-specific; its highest expression in this tissue dominantly correlated with the expression of mitochondrial genes pointing to a possible link between the presence of ThTr2 and brown/beige adipocytes (Pereira et al., 2021). Analysis of RNAseq data of Perdikari et al. (2018) reveals that ThTr2 expression is higher than ThTr1 expression (estimated by mRNA counts) in both human WAT and BAT obtained by needle biopsies from the neck area (Fig. S1c, d). These data along with our observations (Fig.1) suggested that ThTr2 and thiamine might have a role in the regulation of adipocyte thermogenesis.

### Inhibition of ThTr2 hampers uncoupled respiration of adipocytes

To investigate the possible significance of ThTr2 during thermogenic activation in DN and SC derived adipocytes, we treated differentiated adipocytes with dibutyryl-cAMP (which mimics adrenergic stimulation) in the presence of a potent ThTr2 inhibitor, fedratinib (Zhang et al., 2014) and measured OCR in a generally used culture medium which contains thiamine at 8.2 µM concentration. As expected, OCR was elevated immediately upon cAMP addition to DN derived ADIP while the response of SC derived ADIP was moderate. Fedratinib abrogated the cAMP-stimulated elevation of OCR of ADIPs derived from both DN (Fig. 2a, left and middle panels) and SC depots (Fig. 2b, left and middle panels) during the 10 hours monitoring. Proton leak respiration that correlates mainly with UCP1-dependent heat generation was measured upon the injection of the ATP synthase blocker, oligomycin. We found that cAMP-stimulated proton leak respiration was reduced by fedratinib in both DN (Fig. 2a, left and middle panels) and SC (Fig. 2b, left and middle panels) ADIPs. This inhibitory effect of fedratinib was also observed in DN and SC adipocytes with more pronounced browning (B-ADIPs) resulted from their differentiation with constant PPARγ stimulation by rosiglitazone (Fig. S2, left and middle panels).

**Figure 2.**
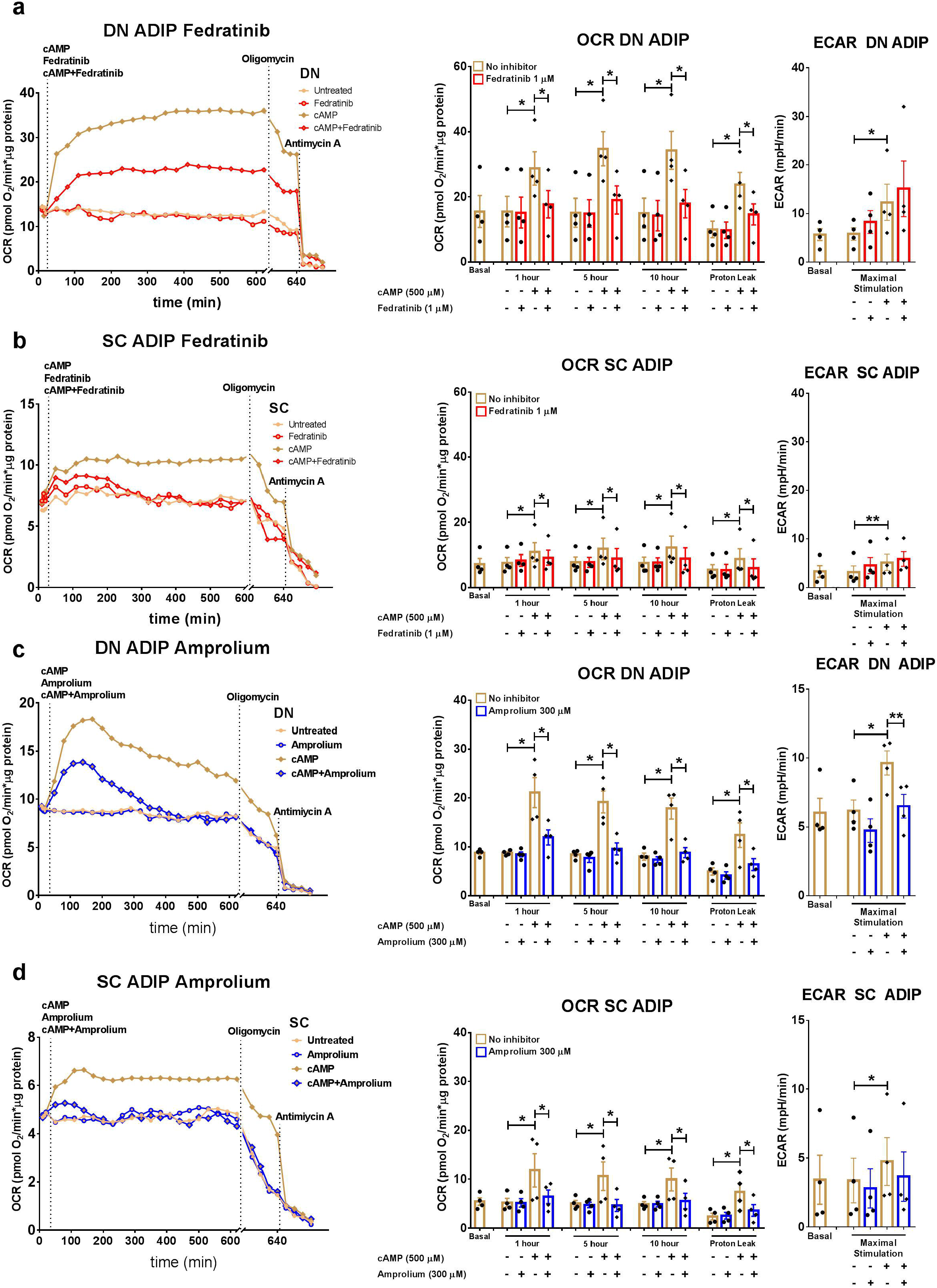
Effect of thiamine transporter inhibitors on the cAMP-induced oxygen consumption (OCR) and extracellular acidification (ECAR) rates in human deep neck (DN) and subcutaneous (SC) derived adipocytes. SC and DN-derived preadipocytes were differentiated into ADIP, then OCRs were detected for 10 hours following the injection of dibutyryl-cAMP in the presence or absence of thiamine transporter inhibitors, fedratinib (a-b) or amprolium (c-d). Each left panel shows representative curves of 4 measurements. OCR at basal, 1, 5, and 10h post-injection, and after oligomycin addition (middle panels), and ECAR (right panels) were quantified in ADIPs derived from four independent donors. Statistical analysis was performed by unpaired t-test, *p<0.05, **p<0.01.

Fedratinib can also inhibit some tyrosine kinases including JAK2 (Zhang et al., 2014) and although tyrosine kinases are not known to be involved in adrenergic signaling of cultured adipocytes, we found it necessary to test also the effect of the thiamine analogue amprolium that inhibits the thiamine transport activity of both ThTr2 and ThTr1, with IC_50_ 0.620±0.27 and 2.60±0.93 µM, respectively (Giacomini et al., 2017). Amprolium was also effective in significantly reducing cAMP-stimulated maximal and proton leak respiration in both DN (Fig. 2c, left and middle panels) and SC (Fig. 2d, left and middle panels) derived ADIPs. cAMP-stimulated ECAR was not affected by the inhibition of ThTr2 by fedratinib in the adipocytes (Fig. 2a-b and Fig S2a-b, right panels) while amprolium treatment led to its decrease (Fig. 2c-d, right panels) in DN ADIP. The inhibitory effects of fedratinib and amprolium on OCR and proton leak respiration clearly show that continuous supply of thiamine through its transporters is needed for efficient thermogenic response of adipocytes.

Oxythiamine is an antivitamin derivative of thiamine which competitively inhibits thiamine transport and also thiamine pyrophosphokinase, restricting the level of TPP, and at the same time become pyrophosphorylated to oxythiamine-TPP which competes with the coenzyme function of TPP (Bunik et al., 2013). We found that oxythiamine decreased cAMP-stimulated OCR of DN (Fig. S3a) and SC ADIPs (Fig. S3b) at 5 and 10 hours after injection. However, proton leak respiration was not affected by oxythiamine in either DN or SC derived ADIPs. ECAR was decreased by oxythiamine in both ADIP types (Fig. S3a-b). It has been reported that oxythiamine does not inhibit ThTrs as efficiently as fedratinib or amprolium (Giacomini et al., 2017) suggesting that high thiamine uptake could still occur in the adipocytes upon oxythiamine treatment which may explain why oxythiamine at the applied concentration did not decrease proton leak respiration.

### Thiamine enhances thermogenic activation of adipocytes in a concentration-dependent manner

As a next step, we studied the importance of thiamine availability during thermogenic activation of adipocytes using a thiamine free culture medium and co-injection of cAMP with gradually increasing concentrations of thiamine (Fig 3a). In the absence of thiamine, as compared to 8.2 µM concentration present in regular culture medium, cAMP-induced OCR to maximal respiration rate was lower in both DN and SC ADIPs (comparing data on Fig 3a and b to Fig 2a and b). cAMP-stimulated maximal respiration was increased already after addition of thiamine at 40 nM concentration and elevated further at rising concentrations up to 25 µM (Fig. 3a) demonstrating that thiamine availability in abundance has critical importance during thermogenic activation of adipocytes. To determine the effect of increasing thiamine concentration on UCP1-dependent portion of cellular respiration, we calculated the OCR observed following oligomycin injection and found that increasing thiamine concentration resulted in higher proton leak respiration which could be observed already at 40 nM thiamine level (Fig. 3a). At this thiamine concentration we estimated the amount of thiamine used up from the culture fluid during 10 hours of cAMP stimulation of DN ADIP derived from four independent donors. By measuring actual thiamine concentrations and comparing culture media of unstimulated and cAMP treated adipocytes, we found that 10^5^ stimulated adipocytes consumed 9.75±3.15 (mean ±SD) pmol thiamine compared to not detectable consumption by unstimulated adipocytes. Thiamine consumption was decreased to 4.32±1.70 and 2.87±0.71pmol (p<0.05) per 10^5^ in the cells presence of 1 µM fedratinib or 300 µM amprolium, respectively.

**Figure 3.**
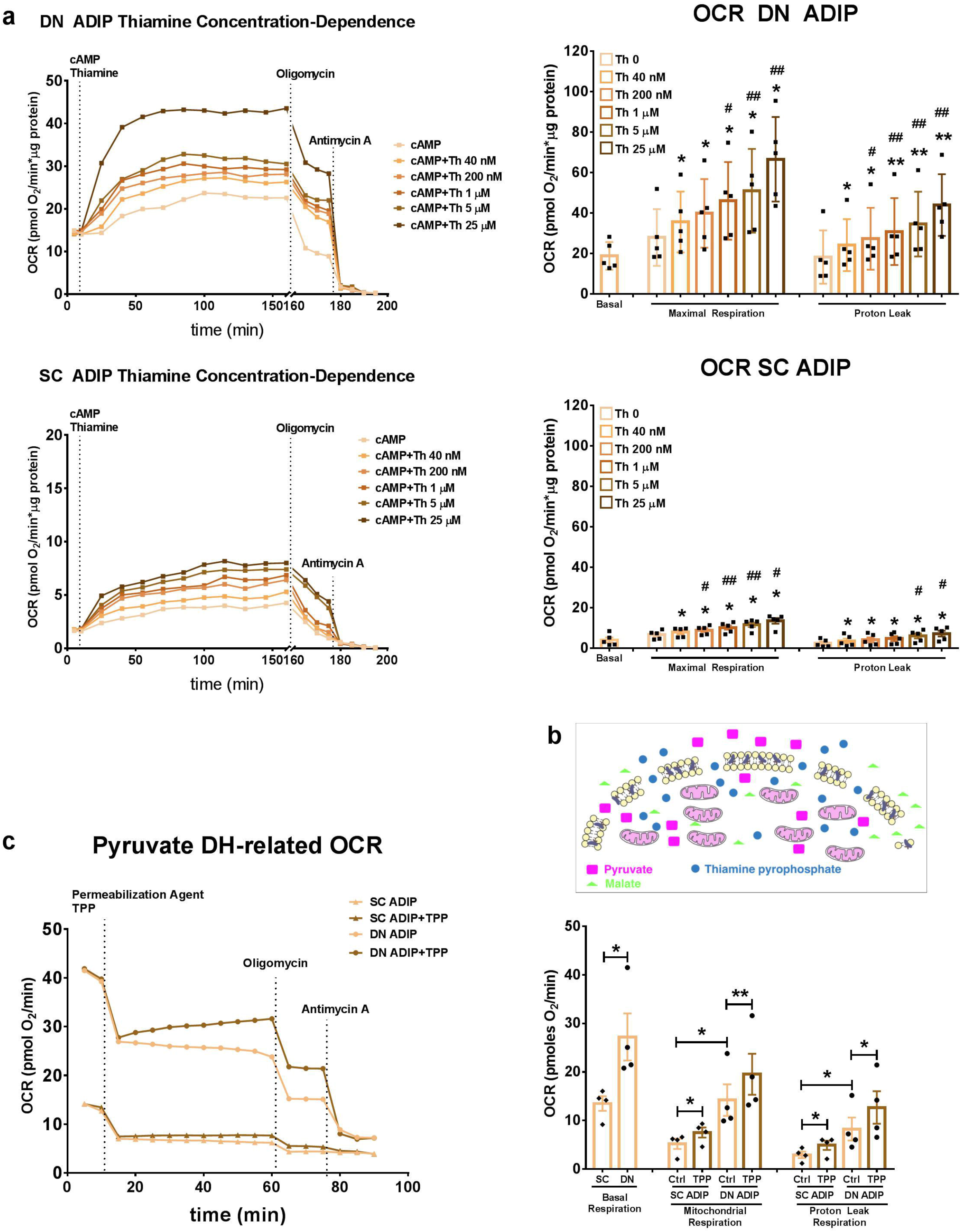
Effect of thiamine (Th) availability and direct stimulation of pyruvate dehydrogenase (PDH) by Th pyrophosphate (TPP) on the thermogenic activation of adipocytes. After differentiation, ADIPs were incubated in Th free media for one hour, then treated with 500 µM dibutyryl-cAMP and gradually increasing concentrations of Th for 10 hours. (a) OCR of DN and SC ADIPs was monitored and representative curves of five measurements (left panels) are shown. OCR at basal and maximal stimulation, and after oligomycin addition (right panels) was quantified in ADIPs derived from five independent donors. Statistical analysis was performed by *comparing effect of each concentration of Th to the lack of Th and #higher than 40 nM concentrations to the effect of 40nM concentration. (b) Scheme of cell membrane permeabilization experiment where ADIPs were incubated in mitochondrial assay solution. OCR was measured following injection of the cell permeabilizer and TPP. (c) Representative curves of four measurements (left panel). OCR at basal, mitochondrial respiration, and proton leak respiration were quantified from four independent donors (right panel). Statistical analysis was performed by unpaired t-test, */#p<0.05, **/##p<0.01.

### Activity of TPP-dependent PDH can be increased by TPP in permeabilized adipocytes

To further understand the mechanism of thermogenic action of thiamine, which is converted to the biochemically active compound TPP in cells, we optimized a Seahorse-based respiration assay to monitor the activity of one of the TPP-dependent enzyme complexes, PDH, in cell membrane-permeabilized adipocytes which enabled us to study mitochondrial function without isolating mitochondria (Fig. 3b). Substrate for PDH and TCA cycle were added to drive NADH generation. Upon the injection of membrane permeabilizer, OCR dropped as expected in both DN and SC derived ADIPs (Fig. 3c). However, the difference in basal OCR between DN and SC ADIPs as well as responsiveness to oligomycin were maintained under the permeabilized conditions. TPP addition significantly increased both maximal mitochondrial and proton leak respiration in DN derived ADIPs while a less pronounced effect was observed in SC ADIP (Fig. 3c). Transport of TPP to mitochondria is mediated by TPC, encoded by *SLC25A19*, which was expressed higher in DN as compared to SC derived adipocytes and cAMP elevated its expression, indicating the increased demand of TPP in the mitochondria during thermogenic activation (Fig. S4a-b). It is worth noting that ThTr2 inhibition did not influence protein expression of TPC or PDH subunit alpha (PDHA1), and the latter was not affected by increasing thiamine availability either (Fig. S4c-d). The results obtained with the permeabilized adipocytes suggest that TPP-dependent PDH entities, which generate metabolic fuel for thermogenesis, are not fully saturated with bound TPP in differentiated adipocytes. This implies that excess thiamine converted to TPP in stimulated adipocytes can increase respiration and thermogenesis through elevating the level of TPP bound enzymes and thereby NADH production.

We also investigated the effect of inhibiting another component of the PDH complex, the E2 subunit, in intact adipocytes. We treated the cells with cAMP analogue in the presence of lipoic acid antagonist, devimistat (Zachar et al., 2011). Lipoic acid is an important cofactor for several mitochondrial enzyme complexes, such as PDH, α-ketoglutarate dehydrogenase, and branched-chain ketoacid dehydrogenase (Solmonson et al., 2018). Devimistat inhibited cAMP-stimulated elevation of OCR in both DN (Fig. S3c) and SC (Fig. S3d) ADIPs at 1, 5, and 10 hours after injection. It increased ECAR in SC but did not affect that in DN ADIP (Fig. S3c-d, right panels). Importantly, proton leak respiration was decreased upon addition of devimistat indicating that continuous availability of co-factors for steady-state activity of TPP-dependent enzymes is critical for maintaining effective thermogenic stimulation.

### Inhibition of thiamine transport leads to lower expression of thermogenic genes in thermogenic adipocytes and adipose tissue biopsies

The hampered thermogenesis observed in the presence of ThTr inhibitors raised the possibility that limited thiamine availability could influence thermogenic gene expression. As described earlier (Toth et al., 2020), DN ADIPs expressed UCP1 (Fig. 4a) and PGC1a (Fig. 4b) at a higher level than SC derived ones. cAMP increased the mRNA and protein expression of these thermogenic marker genes in both SC and DN ADIPs at thiamine concentration present in the standard medium (Fig. 4a-c). Inhibition of ThTrs by either fedratinib or amprolium resulted in attenuated cAMP-dependent upregulation of UCP1 and PGC1a (Fig. 4a-c). Furthermore, the high expression of both UCP1 and PGC1a in DN ADIP was decreased as a result of inhibitor treatments even in the absence of cAMP stimulation suggesting that continuous supply of thiamine is needed to maintain a high basal expression of these thermogenic genes. We observed a reduced expression of additional thermogenic marker genes, such as *DIO2, CITED1, CIDEA*, and *TBX1* in response to fedratinib during thermogenic activation (Fig. 4d-g). Expression of ThTr2 was not affected by the inhibitor suggesting that addition of fedratinib decreased only its activity (Fig. S5a-b). ThTr1 expression was not influenced either (Fig S5c-d).

**Figure 4.**
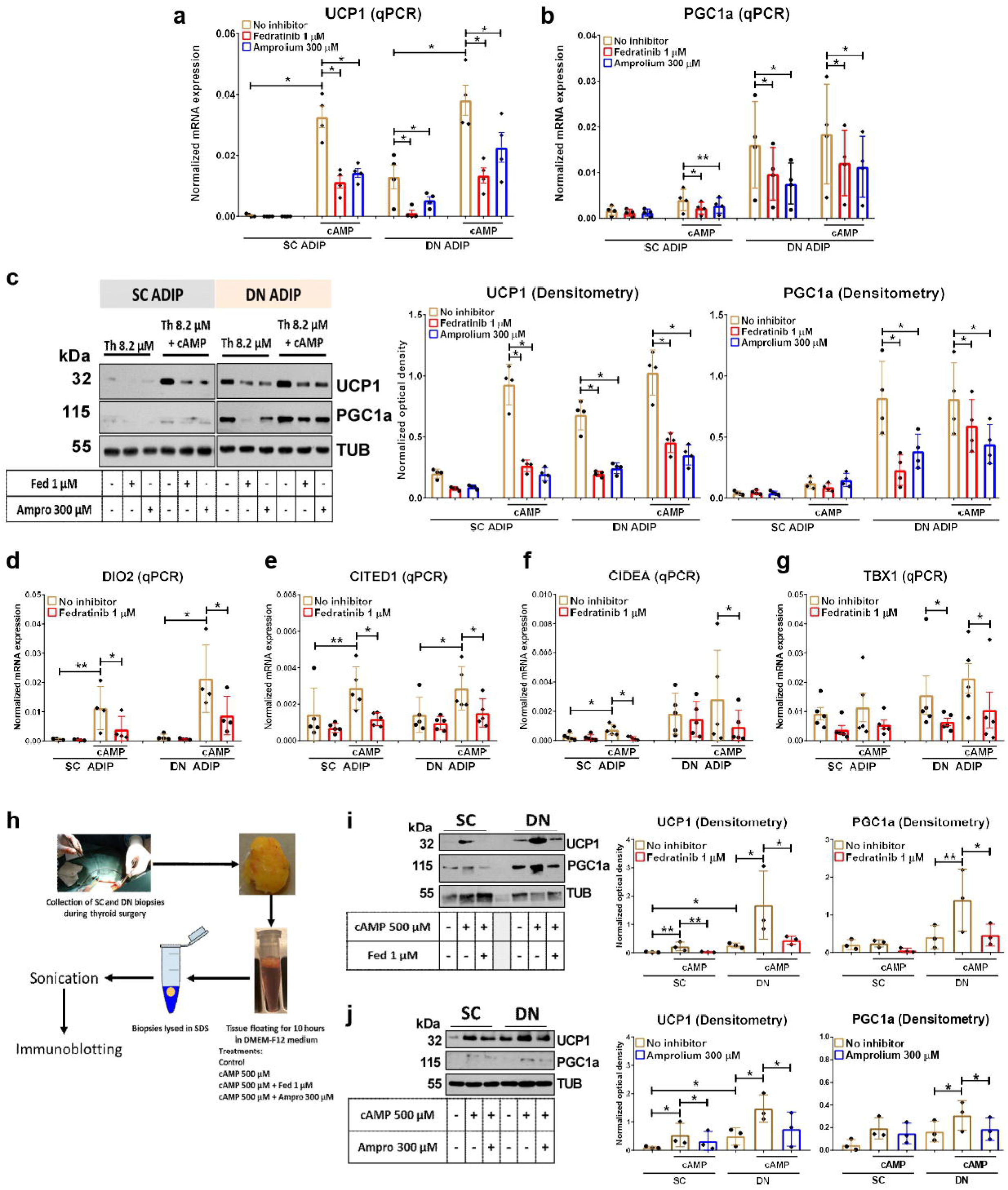
Effect of thiamine transporter inhibitors on the basal and cAMP induced expression of thermogenic marker genes in human deep neck (DN) and subcutaneous (SC) derived adipocytes. After differentiation, ADIPs were treated with 500 µM dibutyryl-cAMP in the presence or absence of thiamine transporter inhibitors, fedratinib (Fed) or amprolium (Ampro) for 10 hours. (a-b) mRNA expression of *UCP1* and *PGC1a* detected by RT-qPCR, n=4. (c) Protein expression of UCP1 and PCG1a detected by immunoblotting, n=4. (d-g) mRNA expression of *DIO2, CITED1, CIDEA*, and *TBX1* detected by RT-qPCR, n=5. (h) Schematic experimental design of human biopsies floating. (i-j) Protein expression of UCP1 and PGC1a, in human SC and DN biopsies detected by immunoblotting in the presence of fed (i) or ampro (j), n=3 (different donors for the fedratinib and amprolium experiments). *p<0.05, **p<0.01, statistical analysis by unpaired t-test.

We also investigated the effect of fedratinib on thermogenic gene expression in SC and DN derived B-ADIPs. We found that cAMP could not elevate further the mRNA and protein expression of UCP1 and PGC1a as their basal expression in B-ADIP was already high (Fig. S6a-c). However, ThTr2 inhibition led to their decreased expression (Fig. S6a-c) and expression of *DIO2* and *TBX1* was also hampered by fedratinib treatment (Fig. S6d,e) demonstrating that thiamine transport is needed for keeping the high expression of thermogenic genes in brown differentiated, thermogenically activated B-ADIPs as well.

The protein expression of mitochondrial complex subunits I-IV showed an increasing trend in cAMP treated ADIPs at the standard level of thiamine in the culture medium and this could be downregulated by either fedratinib or amprolium (Fig. S7a-e). The amount of mitochondrial complex V subunit was not affected by either cAMP or the applied inhibitors (Fig. S7f). Inhibition of TPP requiring enzyme complexes by devimistat resulted in reduced cAMP- and thiamine-dependent upregulation of UCP1 and PGC1a mRNA and protein expression in SC and DN ADIPs (Fig. S8). However, we did not observe any inhibitory effect of oxythiamine on the expression of these genes in ADIPs (Fig. S8).

Next, we addressed the question whether inhibition of ThTrs affects the protein expression of thermogenic markers and ThTrs *in situ*. Pairs of SC and DN biopsies were dissected into three small pieces which were incubated in culture media with 8.2 µM thiamine. The first sample served as a control, the second was subjected to cAMP-mediated thermogenic activation, the third one received co-treatment of cAMP and either fedratinib or amprolium (Fig. 4h). In accordance with the literature and our previous *ex vivo* observations, UCP1 and PGC1a were expressed at a higher level in DN compared to SC biopsies (Fig. 4i, j). UCP1 and PGC1a were remarkably upregulated in response to cAMP-dependent activation, which effect was significantly inhibited by fedratinib and amprolium in both SC and DN adipose tissues (Fig. 4i-j). The expression of ThTrs was not significantly affected by either cAMP or its administration with the ThTrs inhibitors (Fig. S5e-f).

### Thiamine potentiates thermogenic gene induction in adipocytes

As we observed the elevation of proton leak respiration following a gradually increasing concentration of thiamine, we presumed that expression of thermogenic marker genes was also regulated by thiamine availability. Using thiamine free culture fluid, we could observe that cAMP-stimulated upregulation of UCP1 and PGC1a were potentiated by applying increasing concentrations of thiamine (Fig. 5a-c). Regarding UCP1, this effect of thiamine was observed at its 40 nM and 1 µM levels in SC and DN ADIPs, respectively, while PGC1a expression was potentiated at 40 nM in both. We also investigated the effect of thiamine on the mRNA expression of *DIO2, CITED1, CIDEA*, and *TBX1* and found that thiamine potentiated the cAMP induced thermogenic effect in both SC and DN derived ADIPs already at its low concentrations (Fig. 5d-g). These results clearly showed that thiamine could potentiate cAMP induced expression of thermogenic genes.

**Figure 5.**
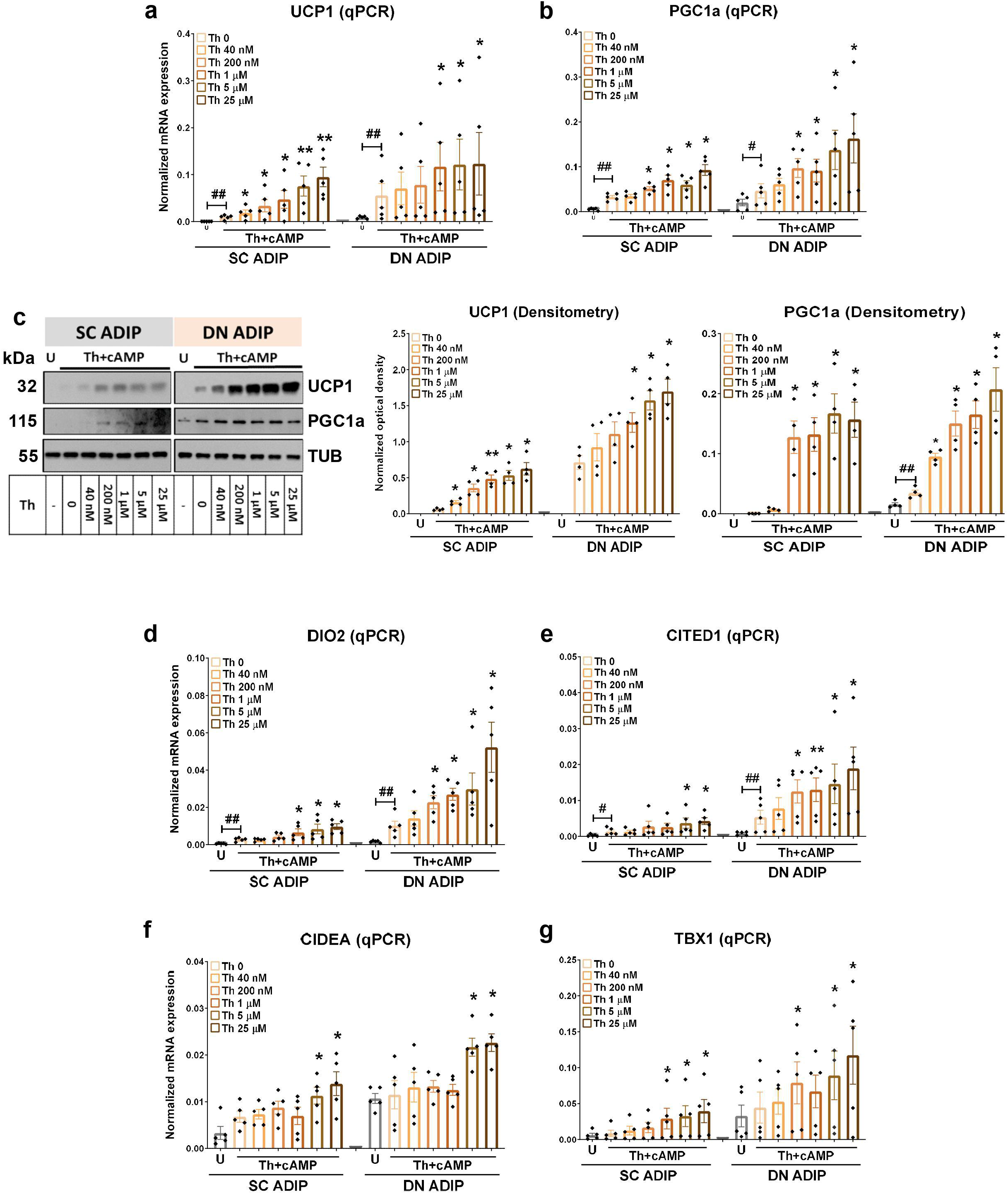
Effect of increasing concentration of thiamine (Th) on thermogenic gene induction in human deep neck (DN) and subcutaneous (SC) derived adipocytes. After differentiation, ADIPs were incubated in Th free media for one hour, then treated with 500 µM dibutyryl-cAMP and gradually increasing concentrations of Th for 10 hours. (a-b) mRNA expression of *UCP1* and *PGC1a* assessed by RT-qPCR, n=5. (c) UCP1 and PGC1a protein expression detected by immunoblotting, n=4. (d-g) mRNA expression of *DIO2, CITED1, CIDEA*, and *TBX1*, n=5. U: untreated. Statistical analysis was performed by unpaired t-test (*#p<0.05, **##p<0.01), *comparing data at each concentration of Th to the lack of Th or # comparing the indicated groups.

Thiamine availability did not affect the mRNA and protein expression of ThTr2 (Fig. S5g, h) in either SC or DN ADIPs. In DN derived ADIPs, we detected increased mitochondrial complex II and IV expression when the adipocytes were treated with thiamine at 1 µM and 200 nM concentrations, respectively, as compared to the lack of thiamine. However, higher concentration of thiamine could not increase the expression of complex II and IV subunits further (Fig. S7). The expression of complex subunits I, III, and V was not affected by thiamine availability (except complex I at high thiamine concentration) in either SC or DN derived ADIPs.

## Discussion

Characterization of brown and beige adipocytes in adipose tissue depots of mice and humans has been significantly advanced during the last couple of years. These cells have to be activated for heat generation, mainly by adrenergic stimuli, and our knowledge about metabolic and regulatory processes of stimulated adipocytes is not complete. According to our presented results, thiamine is essential for adrenergic induction of highly efficient adipocyte thermogenesis. It must be available and transported into adipocytes in sufficient amount to provide TPP for a group of NADH generating enzymes, such as PDH, which lack this essential co-factor, and to potentiate the increased expression of critical thermogenesis genes, such as UCP1.

Physiological thiamine levels in adults are in the nM range varying pursuant to dietary conditions (Whitfield et al., 2018; Gangolf et al., 2010). The concentration of thiamine is much higher in medium of usual cell culture conditions (close to the 10 µM range) ensuring the availability of excess thiamine for cell growth. In case of the studied thermogenic adipocytes, this is sufficiently abundant for maintaining high respiration rate and proton gradient for UCP1 mediated uncoupling upon adrenergic stimulation. The lack of thiamine in the culture fluid almost halted cAMP-stimulated proton leak respiration. Increasing concentration of thiamine generated gradually growing respiration and proton leak, reaching maximum values at µM thiamine levels, which suggests that heat generating capacity of human adipocytes is underutilized at thiamine concentrations present in human tissues (Fig. 3a). Of note, total thiamine concentration in human blood plasma is nearly an order of magnitude lower than it is in rodents and other animals which can easily adapt to cold environmental temperature (Kimura and Itokawa, 1985). In fact, while the total (free + protein-bound) thiamine concentration in human plasma is 10–20 nM in healthy individuals, it is several hundred nM in rodent plasma, which also contains high amounts of thiamine monophosphate (Gangolf et al., 2010; Suzuki et al., 2017). Our finding that human adipocytes require abundant thiamine for high thermogenic response raises the possibility that the low thermogenic activity of human compared to mouse adipose tissues (Sun et al., 2020) can be boosted by significantly increasing the level of thiamine in blood plasma; the latter is achievable with oral consumption of thiamine hydrochloride (Smithline et al., 2012).

Thiamine uptake by cells appears to be mainly mediated by the ThTr1 and 2, which depend on a transcellular proton gradient (Manzetti et al., 2014). Our data show that adipocyte differentiation induced the mRNA expression of ThTr2 and ThTr1 in both SC and DN derived adipocytes, and 10^5^ DN adipocytes consume approximately 10 pmol thiamine during 10 hours of adrenergic stimulation at 40 nM extracellular thiamine concentration. The high affinity ThTr2 (ThTr2 has an influx constant Kt of 25 nM while ThTr1 works at 2.5 μM) (Manzetti et al., 2018) may play a more pronounced role than ThTr1 in the continuous uptake of thiamine during the thermogenic response of adipocytes since its expression level was found to be higher in potentially thermogenic adipocytes (Fig. 1) (Perdikari et al., 2018; Min et al., 2019; Pereira et al., 2021) and its inhibitor fedratinib could significantly decrease respiration in stimulated adipocytes (Fig. 2a-b). We could show the presence of ThTrs at protein level in human SC and DN adipose tissue biopsies where fedratinib and amprolium could inhibit induction of UCP1 and PGC1α by adrenergic stimulation (Fig. 4i-j). These results suggest that the high affinity ThTr2 is available in tissue adipocytes for uptake of high amounts of thiamine, which in turn can facilitate their thermogenesis as we could demonstrate it in the presented experiments using neck area derived adipocytes.

Fedratinib is used in myelofibrosis therapy as a selective inhibitor of JAK2 (Blair, 2019). This raises the possibility that the observed effects of fedratinib on thermogenic activation might be due to inhibition of an activated tyrosine kinase, especially in view that JAK2 KO mice were unable to upregulate BAT UCP1 following a high-fat diet or after cold exposure (Shi et al., 2016). However, involvement of JAK2 in BAT regulation could be demonstrated only *in vivo*, in the presence of non-adipocyte tissue cells (which is not the case in our *ex vivo* differentiated adipocytes) and it was dispensable for induction of UCP1 thermogenesis in WAT through beige adipocytes which are presumed to be dominant among DN adipocytes (Wu et al., 2012; Shinoda et al., 2015; Jespersen et al., 2013; Sanchez-Gurmaches et al., 2016; Toth et al., 2020; Li et al., 2022). Nevertheless, we carried out all of our critical experiments applying the competitive ThTr inhibitor amprolium as well and found that the inhibition of thermogenic response by amprolium was comparable to the fedratinib effect (Fig. 2c-d). We also observed the inhibitory effect of both fedratinib and amprolium on thiamine consumption by DN derived adipocytes during their adrenergic stimulation.

Cells with high demand of metabolic fuel, such as active brown/beige adipocytes, need extra amount of thiamine to boost the activity of TPP-requiring pyruvate and α-ketoglutarate dehydrogenases that generate reduced equivalents for mitochondrial proton gradient. It is generally assumed that in adipocytes these enzymes are fully saturated with TPP at the end of adipocyte differentiation taking place at high thiamine levels which can provide sufficient TPP for turnover of the enzymes and its prosthetic groups. Seeing the strong stimulatory effect of thiamine on maximal and proton leak respiration of adipocytes activated for thermogenesis, we speculated that part of the TPP-requiring, NADH generating enzyme population does not contain TPP before thermogenic activation. Indeed, we could observe direct stimulation of respiration after addition of TPP to cell membrane-permeabilized adipocytes (differentiated in thiamine rich culture fluid) under condition optimized for monitoring PDH-dependent NADH generation (Fig. 3c), and where mitochondrial pyruvate carrier-1 (MPC1) could mediate the transport of pyruvate from the cytosol to mitochondria to fuel the TCA cycle (Panic et al., 2020). The protein expression of the TPP binding PDHA1 was not affected either by ThTr inhibitors or thiamine availability in intact adipocytes (Fig. S4c, d). Therefore, it is likely that thermogenic activation of adipocytes in the presence of abundant thiamine results in increased number of TPP containing PDH (and likely α-ketoglutarate dehydrogenase) subunits which may explain how transported thiamine can increase maximal and uncoupled respiration. Further studies are needed to clarify the mechanism how adipocytes regulate TPP availability for TCA cycle enzymes during thermogenesis. A possible explanation is that they increase the activity of the mitochondrial TPP transporter, TPC. Significantly elevated TPC mRNA was observed in human BAT as compared to WAT (Perdikari et al., 2018) (Fig. S1e). We also found that the mRNA and protein expression of TPC was elevated upon cAMP treatment of DN derived ADIPs (Fig. S4a, b) suggesting that TPC may provide the extra TPP needed in the mitochondria during thermogenic activation for higher PDH and α-ketoglutarate dehydrogenase activity of the TCA cycle. TPP stimulation of mitochondrial respiration of differentiated cells is not unique for adipocytes: a study using mitochondria isolated from mouse brain showed that TPP treatment elevated OCR and ATP turnover (Ikeda et al., 2016).

It was a surprising finding in our study that thiamine availability could be linked to regulation of thermogenic gene expression. Inhibition of thiamine transport into stimulated adipocytes resulted in lower the expression of UCP1 and other thermogenic markers induced upon adrenergic stimulation (Fig. 4a-g). Addition of thiamine in thiamine free culture condition to the stimulated adipocytes could increase the expression of these genes in a concentration dependent manner (Fig. 5). We can only speculate about the mechanism of how thiamine may contribute to induction of thermogenic genes during adrenergic stimulation of adipocytes. The 5 non-coding region of the UCP1 gene contains regulatory elements that confer tissue specificity, differentiation dependence, and neuro-hormonal regulation to UCP1 gene transcription (Villaroya et al., 2017) through interactions with a large number of transcription regulators including cAMP-responsive transcription factors as well as the PGC-1a co-regulator. Transcriptional regulation of the UCP1 gene by cAMP-mediated signaling is provided by protein kinase A mediated rapid phosphorylation of the CREB transcription factor at the proximal UCP1 promoter region and p38 MAP kinase-mediated phosphorylation of ATF2 at the upstream enhancer region (Robidoux et al., 2005) constituting a fast mechanism of regulation. Thiamine or TPP may directly influence this concerted transcriptional regulation of UCP1 and other thermogenic genes at these complex regulatory sites. Such an effect of thiamine on gene transcription was reported earlier observing inhibition of p53 DNA binding by thiamine in living cells (McLure et al., 2004). Phosphorylation mediated or TPP-dependent metabolic changes occurring during thermogenesis may also generate so far not identified, indirect gene regulatory signals.

Thiamine deficiency leads to several neurological diseases, such as Beriberi or Wernicke’s encephalopathy and Korsakoff psychosis referred to as Wernicke-Korsakoff syndrome (WKS) (Cook et al., 1998). The high sensitivity of humans to thiamine deficiency is probably linked to low circulating thiamine concentrations and low TPP tissue content (Gangolf et al., 2010). The main symptoms of WKS are ataxia, memory impairment, confabulation, and hypothermia which is often reported as a secondary symptom. One of the main causes of these diseases is chronic alcohol consumption which leads to impaired thiamine absorption. It has been reported that thiamine deficiency leads to the lesion of the hypothalamus, which plays an important role in regulating body temperature, appetite, and vegetative functions (Tanev et al., 2008). Human case studies have reported that the hypothermic condition can be restored after 2 days of parenteral administration of thiamine (Hansen et al., 1984). Our results suggest that in addition to causing impairment of hypothalamic thermoregulation, thiamine deficiency may also compromise peripheral thermogenesis in brown/beige adipocytes contributing to hypothermia of WKS patients.

Alcohol consumption also positively correlates with visceral fat accumulation in healthy individuals (Kim et al., 2012; Dorn et al., 2003), which may be partially caused by the disturbance of thiamine absorption and metabolism. Thiamine deficiency has also been reported in patients with type 1 or 2 diabetes (Thornalley et al., 2007) and thiamine supplementation (100 mg, 3×100 mg daily which is about 100 times higher than the recommended daily allowance (Maguire et al., 2018)) for 6 weeks improved glucose tolerance in hyperglycemic individuals (Alaei et al., 2013). It has also been reported that thiamine supplementation may provide beneficial effects in patients with type 2 diabetes by improving lipid and creatinine profiles (Al-Attas et al., 2014). Individuals with obesity exerted significant thiamine deficiency before undergoing bariatric surgery (Carrodeguas et al., 2005; Flancbaum et al., 2006; Nath et al., 2017; Peterson et al., 2016; Aron-Wisnewsky et al., 2016). A study in Thailand revealed a 42% prevalence of thiamine deficiency among children with obesity (Densupsoontorn et al., 2019). Another study in Mexican American children show that low thiamine intake is inversely associated with higher adiposity (Gunanti et al., 2014). Our previous study revealed that ThTr2 expression was lower in neck fat derived adipocytes isolated from individuals carrying FTO rs1421085 obesity risk alleles CC compared to risk-free alleles TT (Toth et al., 2020). A recent report demonstrated that SC adipose tissue isolated from individuals with obesity expressed lower ThTr2 as compared to lean individuals and the expression of ThTr2 positively correlated with weight loss (Pereira et al., 2021) suggesting its possible role in augmentation of energy metabolism. Thiamine supplementation prevented obesity and obesity-associated metabolic disorders in OLETF rats (Tanaka et al., 2010). Altogether, these reports and our presented findings on the stimulatory effect of thiamine on thermogenesis evoke the possibility of using thiamine in future anti-obesity protocols to augment heat generation by thermogenic adipocytes.

## Methods

## Materials

All chemicals were from Sigma-Aldrich (Munich, Germany) unless stated otherwise.

### Ethics statement and obtained tissue samples

Tissue collection was approved by the Medical Research Council of Hungary (20571-2/2017/EKU) followed by the EU Member States’ Directive 2004/23/EC on presumed consent practice for tissue collection. All experiments were carried out in accordance with the guidelines of the Helsinki Declaration. Written informed consent was obtained from all participants before the surgical procedure. During thyroid surgeries, a pair of DN and SC adipose tissue samples was obtained to rule out inter-individual variations. Patients with known diabetes, malignant tumor or with abnormal thyroid hormone levels at the time of surgery were excluded. Human adipose-derived stromal cells (hASCs) were isolated from SC and DN fat biopsies as described previously (Toth et al., 2020; Kristof et al., 2019). Wherever indicated, 10-30 mg of SC and DN tissue samples from the same donors were floated for 10h in DMEM-F12-HAM, at 37 ^0^C and 5% CO_2_ in empty medium, in the presence of 500 µM dibutyryl-cAMP (cat#D0627) alone or in the presence of 500 µM dibutyryl-cAMP and 1µM fedratinib (Selleck Chemicals LLC cat#S2736, Houston, TX, USA) or amprolium 300 µM (cat#137-88-2) (Kristof et al., 2019).

### Differentiation and treatment of hASCs

Human primary adipocytes (ADIP) were differentiated from SVF of adipose tissue containing preadipocytes according to a described protocol applying insulin, cortisol, T3, dexamethasone and short term rosiglitazone treatment (Kristof et al., 2015). We also differentiated preadipocytes under long-term rosiglitazone effect resulting in higher browning capacity of the adipocytes (B-ADIP) (Toth et al., 2020; Elabd et al., 2009). In DMEM-F12-HAM medium, ADIP and B-ADIP were treated with a single bolus of 500 µM dibutyryl-cAMP for 10 hours to mimic *in vivo* cold-induced thermogenesis (Arianti et al., 2021). Fedratinib 1µM or amprolium 300 µM was administered to inhibit ThTr activity. Oxythiamine 300 µM (cat#614-05-1) or devimistat 75 µM (MedChem Express cat#HY-15453, Monmouth Junction, NJ, USA) was administered to inhibit the activity of TPP-requiring enzymes. In thiamine concentration dependence thermogenic experiments, we used custom-made thiamine free culture fluid (Gibco cat#ME2107367, Waltham, MA, USA) treating adipocytes with the gradually increasing concentration of thiamine (cat# 731188), 40 nM, 200 nM, 1 µM, 5 µM, or 25 µM in the presence or absence of cAMP stimulation for 10 hours.

### RNA isolation and quantitative real time PCR (RT-qPCR)

Cells were collected, total RNA was isolated, and RT-qPCR was performed as described previously (Klusoczki et al., 2019; Kristof et al., 2016). Gene primers and probes were designed and supplied by Thermo Fisher Scientific (Waltham, MA, USA) as listed in Supplementary Table 1.

### Immunoblotting and densitometry

Immunoblotting and densitometry were carried out as described previously (Szatmari-Toth et al., 2020). Antibodies and working dilutions are listed in Supplementary Table 2.

### Oxygen consumption (OCR) and extracellular acidification rate (ECAR) measurement

OCR and ECAR of adipocytes were measured using an XF96 oxymeter (Seahorse Biosciences, North Billerica, MA, USA) as described previously (Arianti et al., 2021; Kristof et al., 2016). After recording the baseline OCR, 500 µM dibutyryl-cAMP, 1µM fedratinib, 300 µM amprolium, 300 µM oxythiamine, 75 µM devimistat, or combination of cAMP and one of the inhibitors were injected to the cells.

For thiamine concentration dependence experiment, we incubated the adipocytes in thiamine free culture fluid 1h before the extracellular flux assay. After recording the baseline OCR, 500 µM dibutyryl-cAMP and gradually increasing concentrations of thiamine were co-injected. Then, stimulated OCR was recorded every 30 minutes in both types of experimental settings. Proton leak respiration was monitored after injecting oligomycin (cat#495455) at 2□μM concentration. Cells received a single bolus of Antimycin A (cat#A8674) at 10□μM concentration for baseline correction (measuring non-mitochondrial respiration). The OCR was normalized to protein content.

To study the direct effect of TPP on PDH activity, a Seahorse-based assay was optimized for adipocytes (Mikulas et al., 2020). To SC and DN derived adipocytes a mitochondrial assay solution containing 600 mM mannitol (cat#M4125), 210 mM sucrose, 30 mM KH_2_PO_4_, 1.5 mM MgCl_2_, 6mM HEPES, 3 mM EGTA, 0.6% BSA, 4 mM ADP (cat#01905), and 10 mM pyruvate (cat#P76209) + 5 mM malate (cat#M7397), pH 7.4, was added before the measurement. Membrane permeabilizer (Agilent cat#102504-100, Santa Clara, CA, USA) at 1 nM concentration and 300 µM TPP (cat#PHR1369) were administered to the adipocytes after recording the basal OCR for 10 minutes. Then, OCR of permabilized adipocytes was recorded every 5 minutes. Proton leak respiration was determined after injecting oligomycin at 2□μM concentration. Permeabilized cells received a single bolus of Antimycin A at 10□μM concentration for baseline correction.

### Measurement of thiamine concentration

The cell culture fluids were collected, then filtered using Nanosep 3 kDa spin column (Pall Corp, New York, NY, USA) at 12,800x g, 4 ^o^C for 10 min, to get rid of the debris and the high molecular weight components. The filtrates were spiked with stable isotope-labeled (SIL) thiamine (cat#731188) in 2 ng/ml final concentration and introduced to the autosampler of the UPLC. 1 μl of the sample was injected in duplicates and the separation was done on H-class UPLC (Waters, Milford, MA, USA) using a 17.5-minute gradient and AccQ-TagTM ULTRA C18 1.7 μm, 2.1 × 100 mm (Waters, Milford, MA, USA) analytical column. Buffer A was 10 mM ammonium formate, 0.1% formic acid in water, the buffer B was 10 mM ammonium formate, 0.1% formic acid in methanol. The UPLC was coupled online to a 5500 QTRAP (Sciex, Framingham, MA, USA) mass spectrometer and the SRM transitions characteristic for SIL thiamine (268.8/148, 268.8/122, 268.8/81) and endogenous thiamine (265/144, 265/122, 265/81) were recorded. The area under the curve (AUC) values were calculated for the transitions and the compound-specific values were summed in the case of each compound. Normalization was carried out using the mean AUC values for SIL thiamine and the normalized values were used to calculate the thiamine concentration by utilizing a 7-point calibration curve in the range of 0.5-40 ng/ml of thiamine.

## Statistical analysis

The results are expressed as mean±SD. Normality of distribution of the data was tested by Kolmogorov-Smirnov test. In comparison of two groups, unpaired t-test was used. The data were visualized and analyzed by using GraphPad Prism 8 (GraphPad Software, San Diego, CA, USA).

## Author Contribution

LF and RA conceived and designed the experiments with contributions from EK. RA, BÁV and EK performed the experiments. FG provided tissue samples. AG and EC measured thiamine concentrations in culture fluid. RA and LF wrote the manuscript with inputs from EK.

## Acknowledgement

We thank Dr. Beáta B. Tóth for her exceptional help in data analysis and reviewing the manuscript before its submission, László Tretter (Semmelweis University, Budapest) for advising in the design of cell permeability experiments, Abhirup Shaw and Attila Vámos for the help in cell preparation, and Jennifer Nagy for technical assistance. This research was funded by the European Union and the European Regional Development Fund (GINOP-2.3.2-15-2016-00006) and the National Research, Development and Innovation Office (NKFIH-FK131424 and K129139) of Hungary. EK was supported by the János Bolyai Fellowship of the Hungarian Academy of Sciences. RA was supported by Stipendium Hungaricum and Indonesian Education Scholarship (LPDP Indonesia). BÁV was supported by the ÚNKP-21-3-I New National Excellence Program of the Ministry for Innovation and Technology from the source of the National Research, Development and Innovation Fund.

