## Supplementary Figures and Tables for "Availability of abundant thiamine determines efficiency of thermogenic activation in human neck area derived adipocytes"

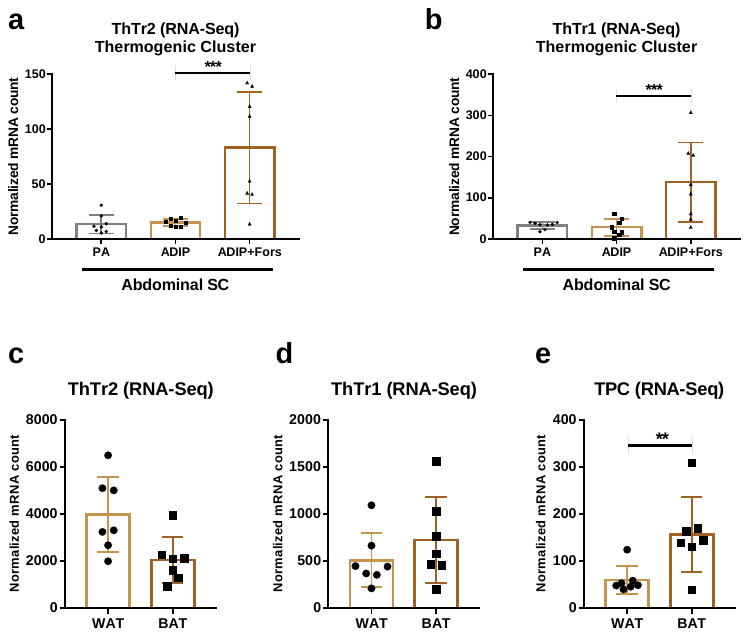

Figure S1. The expression of thiamine transporters (ThTrs) in human abdominal subcutaneous (SC) adipocytes (ADIP) and neck tissue. (a-b) Normalized mRNA counts of ThTr2 and ThTr1 of thermogenic cluster of human abdominal SC derived ADIP (n=8). Based on available RNA-Seq data ([www.ncbi.nlm.nih.gov/geo](http://www.ncbi.nlm.nih.gov/geo), accession number GSE134570 (Min et at., 2019)). PA: preadipocytes; Fors: forskolin. (c-e) Normalized mRNA counts of ThTr2, ThTr1, and the mitochondrial TPP transporter (TPC) of human white and brown adipose tissue (WAT and BAT) obtained by needle biopsies. Based on available RNA-Seq data retrieved from the European Nucleotide Archive, accession number PRJEB20634 (Perdikari et al., 2018). n=7. Statistical analysis of FASTQ files aligned to BAM were performed by DESeq2. **p<0.01, ***p<0.0001.

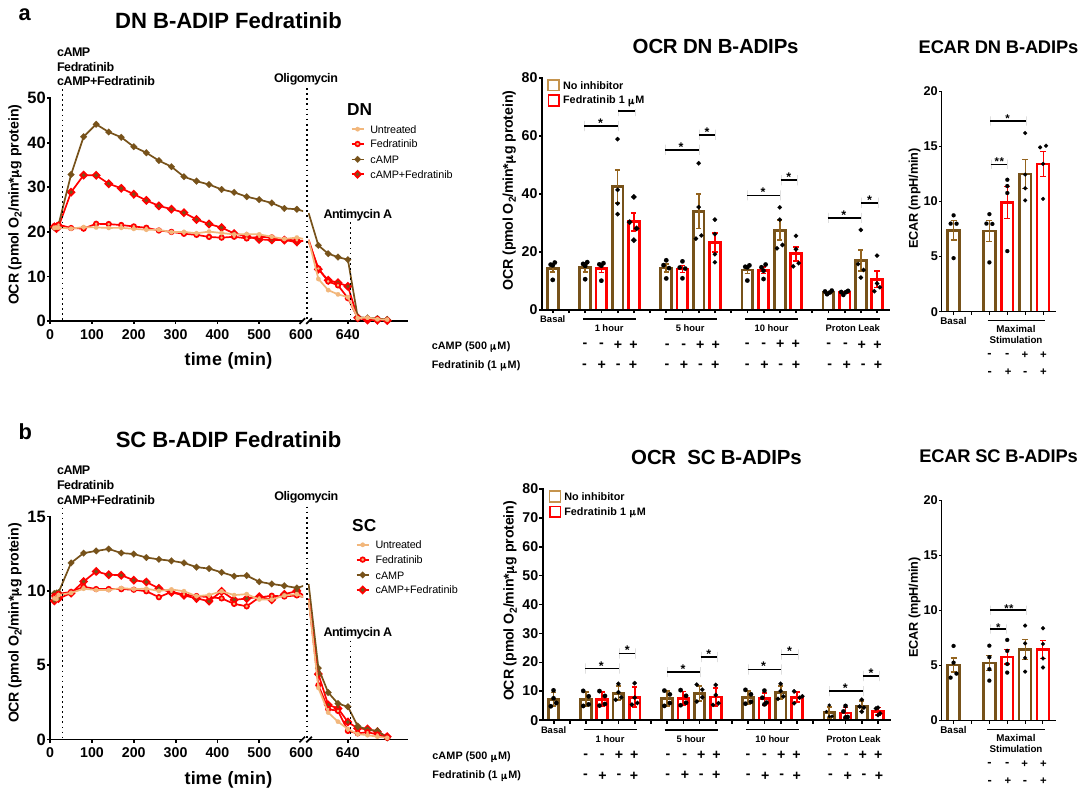

Figure S2. Effect of thiamine transporter 2 inhibitor (fedratinib) on oxygen consumption (OCR) and extracellular acidification (ECAR) rates in brown differentiated (B-ADIP) deep neck (DN) and subcutaneous (SC) derived adipocytes. SC and DN progenitors were differentiated into brown adipocytes (B-ADIP) under long-term rosiglitazone treatment. After differentiation, B-ADIPs were treated with cAMP in the presence or absence of fedratinib. OCR in DN (a) and SC (b) B-ADIPs was detected for 10 hours; representative curves of four measurements (left panels). OCR at basal, maximal stimulation, and after oligomycin addition (middle panels) and ECAR (right panels) were quantified in B-ADIPs derived from four independent donors. Statistical analysis was performed by paired t-test, *p<0.05, **p<0.01.

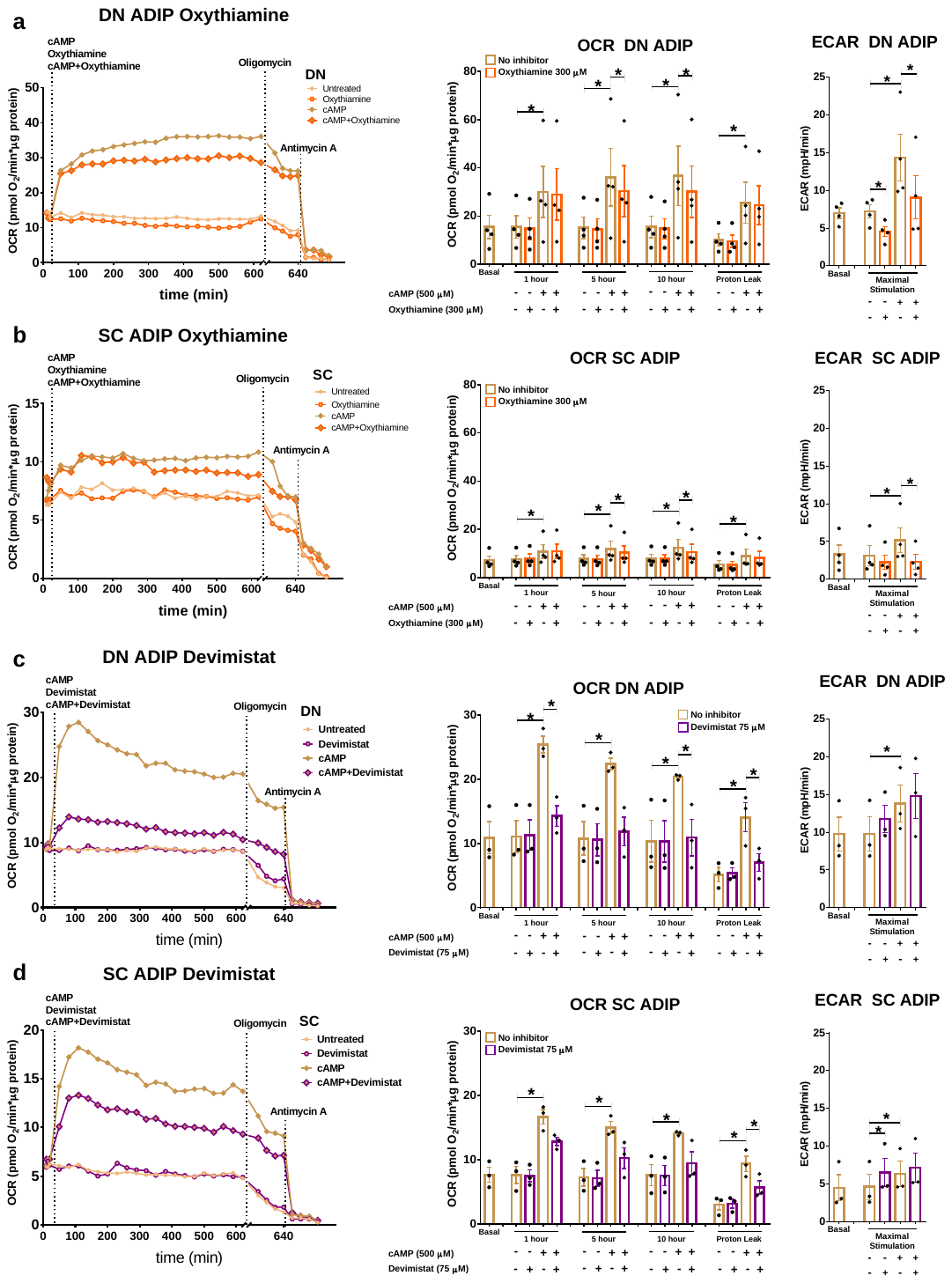

Figure S3. Effect of TPP-dependent enzyme inhibitors on the cAMP-induced expression of oxygen consumption (OCR) and extracellular acidification (ECAR) rates in human deep neck (DN) and subcutaneous (SC) derived adipocytes. ADIPs were differentiated as in Figure 2; OCR was detected for 10 hours following the injection of dibutyryl-cAMP in the presence or absence of oxythiamine (a-b) or devimistat (c-d); representative curves of three or four measurements (left panels). OCR at basal, 1, 5, and 10 hours post-injection, and after oligomycin addition (middle panels), and ECAR (right panels) were quantified in ADIPs derived from three or four independent donors. Statistical analysis was performed by paired t-test, *p<0.05.

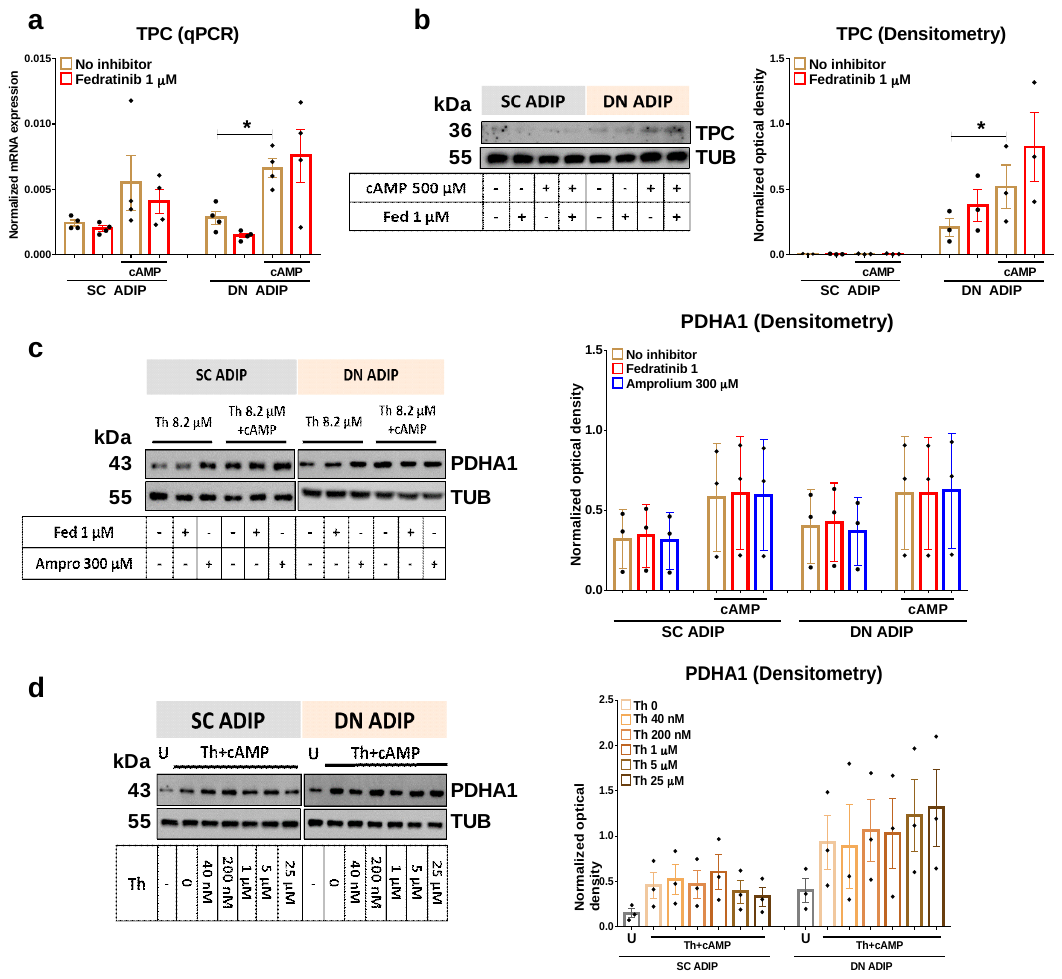

Figure S4. The expression of mitochondrial TPP transporter (TPC) and pyruvate dehydrogenase subunit alpha (PDHA1) in human deep neck (DN) and subcutaneous (SC) derived adipocytes (ADIP). (a) mRNA (n=4) and (b) protein expression (n=3) of TPC in SC and DN derived ADIPs treated with 500 µM dibutyryl-cAMP in the presence of fedratinib (Fed) for 10 hours. (c) Protein expression of PDHA1 in SC and DN derived ADIPs treated with 500 µM dibutyryl-cAMP in the presence of fedratinib or amprolium (Ampro) for 10 hours, n=3. (d) Protein expression of PDHA1 in SC and DN derived ADIPs treated with 500 µM dibutyryl-cAMP and gradually increasing concentrations of thiamine (Th) for 10 hours, n=3. U=untreated. Statistical analysis was performed by paired t-test. *p<0.05.

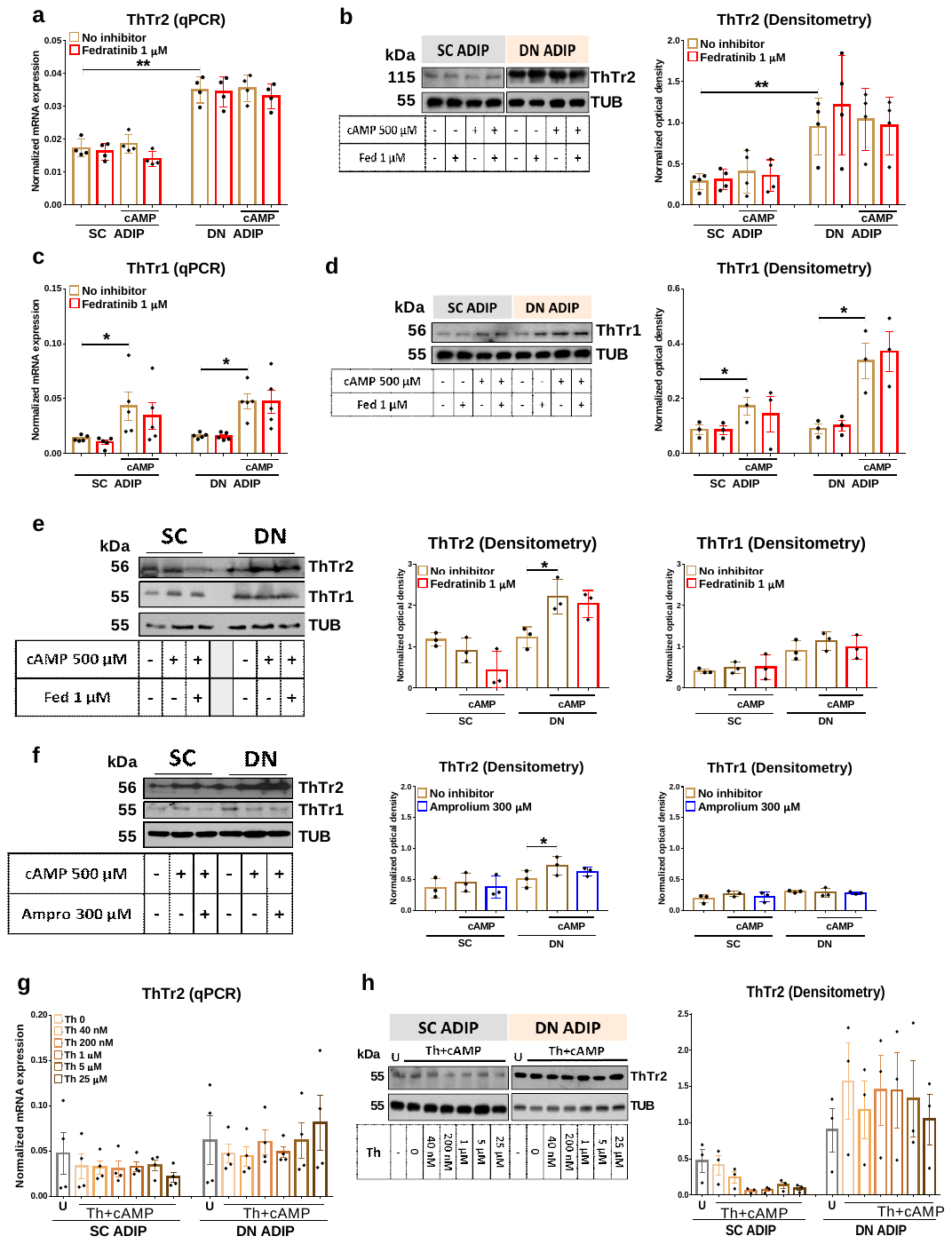

Figure S5. The expression of ThTr2 and ThTr1 was not affected by fedratinib (Fed) in human deep neck (DN) and subcutaneous (SC) adipocytes. ADIPs were differentiated and treated as in Figure 4. After differentiation, ADIPs were treated with 500 µM dibutyryl-cAMP in the presence or absence of Fed for 10 hours. (a) mRNA expression of ThTr2 detected by RT-qPCR, n=4. (b) Protein expression of ThTr2 detected by immunoblotting, n=4. (c) mRNA expression of ThTr1 detected by RT-qPCR, n=4. (d) Protein expression of ThTr1 detected by immunoblotting, n=3. (e-f) Protein expression of ThTr2 and ThTr1 in tissue biopsies treated with cAMP or combination of cAMP and Fed (e) or amprolium (Ampro) (f) for 10 hours, n=3. (g-h) mRNA and protein expression of ThTr2 of ADIPs incubated in thiamine free media for one hour, then treated with 500 µM dibutyryl-cAMP and gradually increasing concentrations of thiamine for 10 hours. Statistical analysis was performed by paired t-test, *p<0.05, **p<0.01.

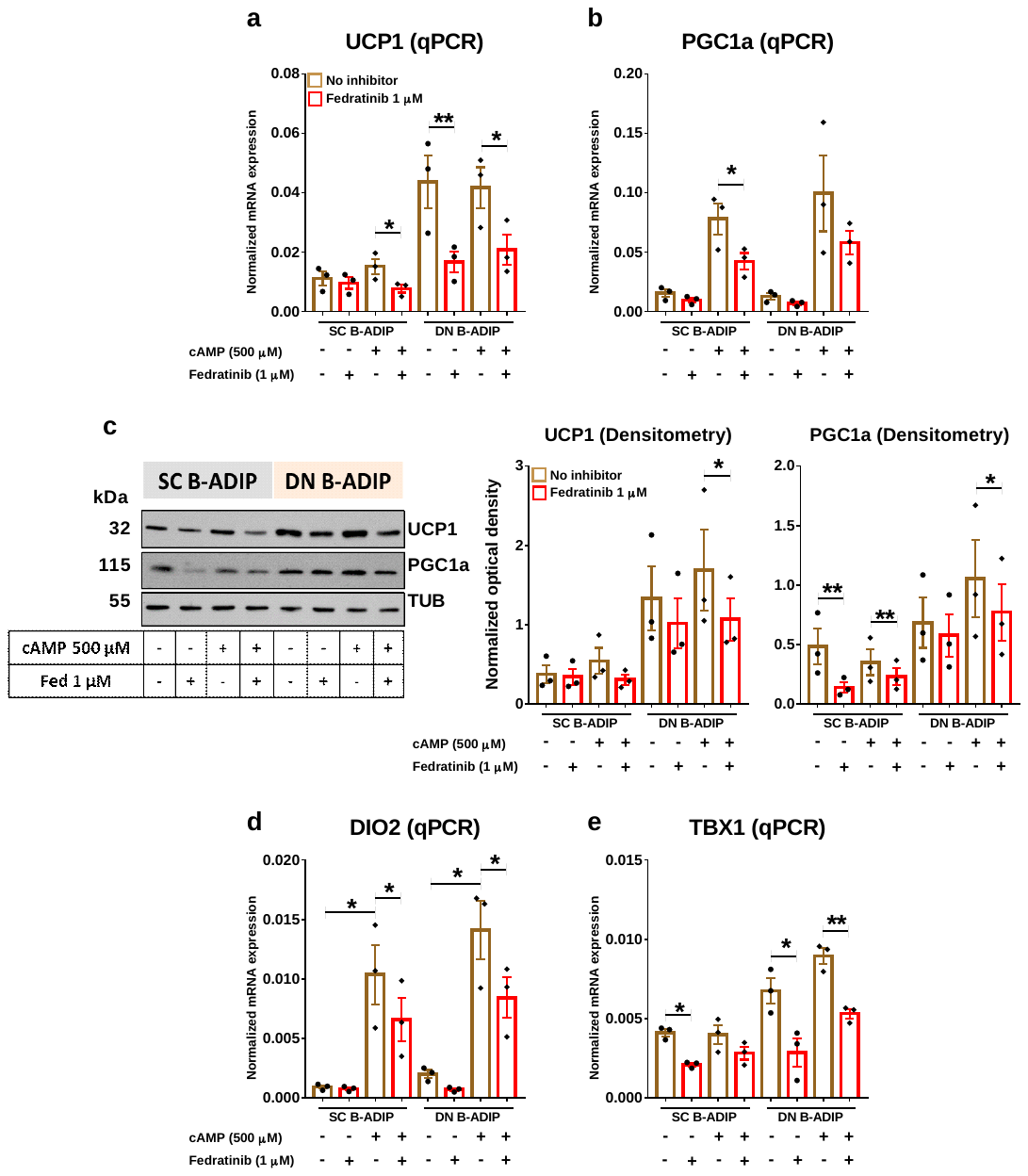

**Figure S6.** Effect of thiamine transporter 2 inhibitor fedratinib (Fed) on the expression of thermogenic markers of subcutaneous (SC) and deep neck (DN) adipocytes differentiated with long term rosiglitazone treatment (B-ADIP). After differentiation, B-ADIPs were treated with 500 µM dibutyryyl-cAMP in the presence or absence of Fed for 10 hours. (a-b) mRNA expression of *UCP1* and *PGC1a* detected by RT-qPCR. (c) Immunoblotting and densitometry of UCP1 and PGC1a protein expression in SC and DN B-ADIPs. (d, e) mRNA expression of *DIO2* and *TBX1* detected by RT-qPCR. n=3 for all groups. Statistical analysis was performed by paired t-test, *p<0.05, **p<0.01.

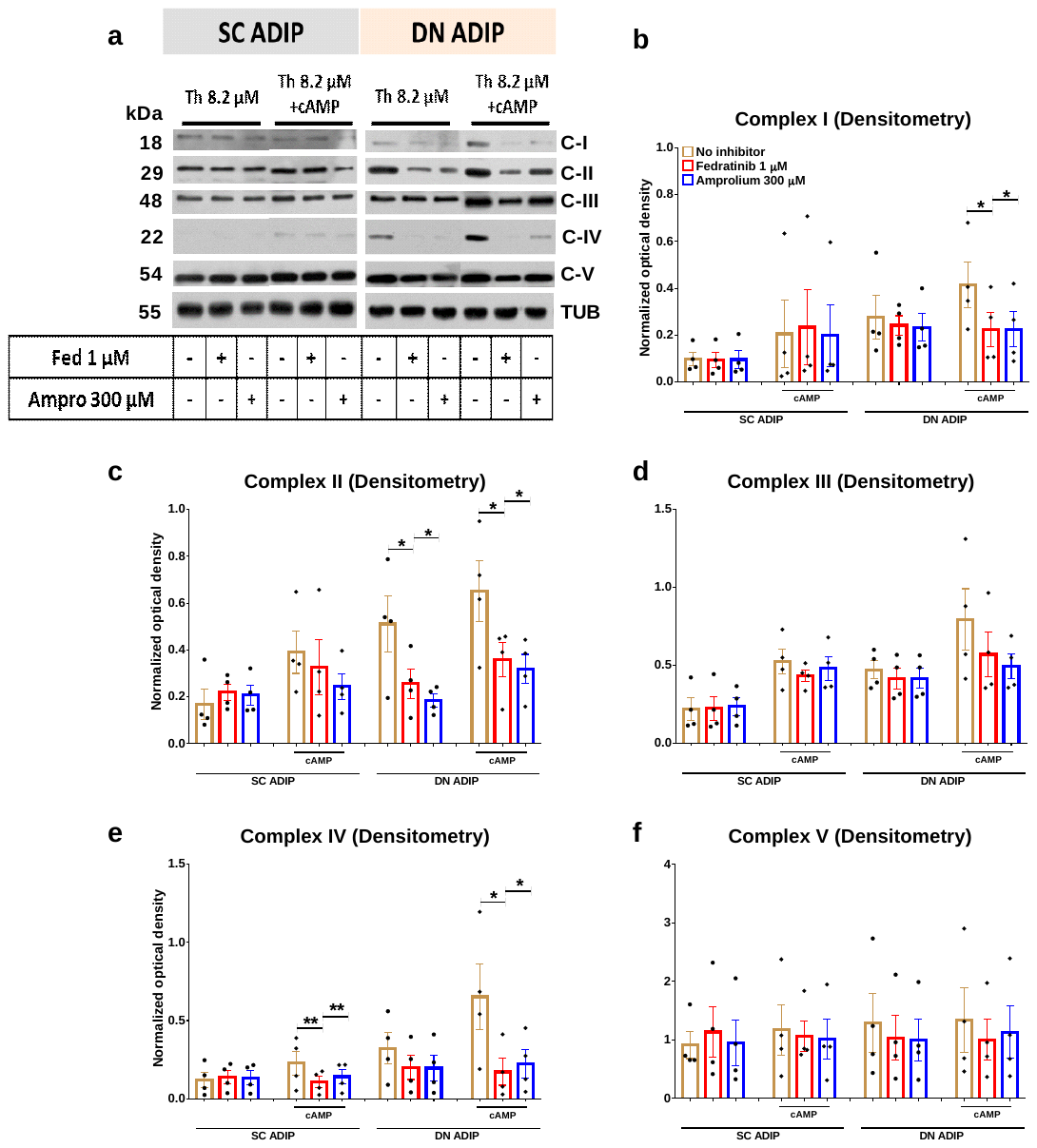

Figure S7. Effect of thiamine (Th) transporter inhibitors on the protein expression of mitochondrial complex subunits in human deep neck (DN) and subcutaneous (SC) derived adipocytes. ADIPs were differentiated and treated as in Figure 4. After differentiation, ADIPs were treated with 500 µM dibutyryl-cAMP in the presence or absence of fedratinib (Fed) or amprolium (Ampro) for 10 hours. (a) Expression of mitochondrial complex subunits detected by immunoblotting. (b-f) Quantification of complex I-V immunoblotting by densitometry; n=3. Statistical analysis was performed by paired t-test, *p<0.05, **p<0.01.

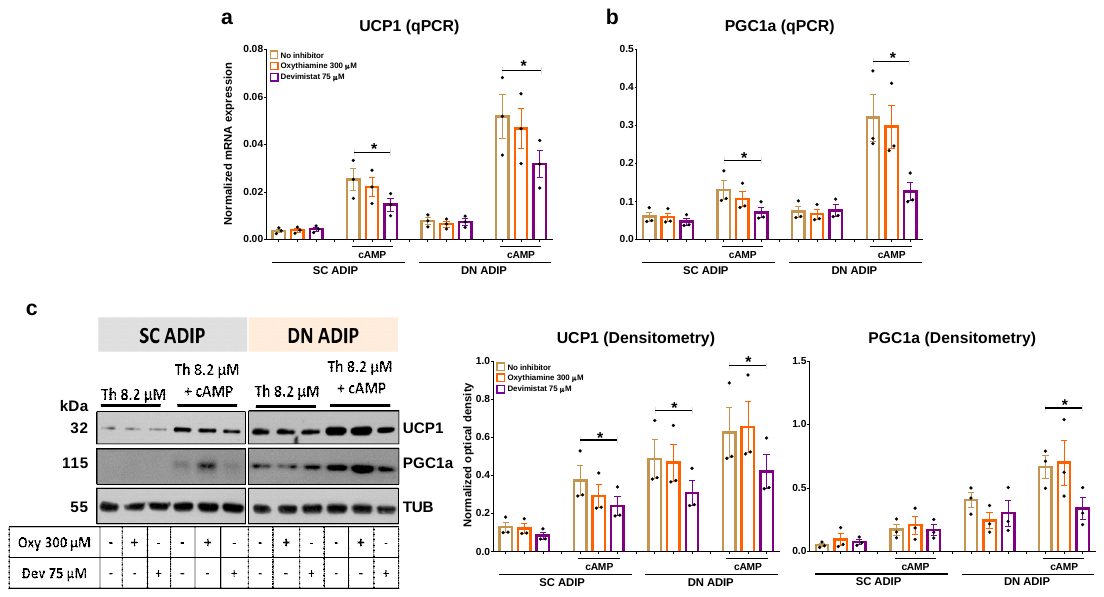

Figure S8. Effect of oxythiamine (Oxy) and devimistat (Dev) on the expression of thermogenic marker genes in human deep neck (DN) and subcutaneous (SC) derived adipocytes. ADIPs were differentiated and treated as in Figure 4. After differentiation, ADIPs were treated with 500 µM dibutyryl-cAMP in the presence or absence of Oxy or Dev for 10 hours. (a-b) mRNA expression of *UCP1* and *PGC1a* detected by qPCR. (c) Immunoblotting and densitometry of UCP1 and PGC1a protein expression in SC and DN ADIPs. n=3 for all groups. Statistical analysis was performed by paired t test, *p<0.05.

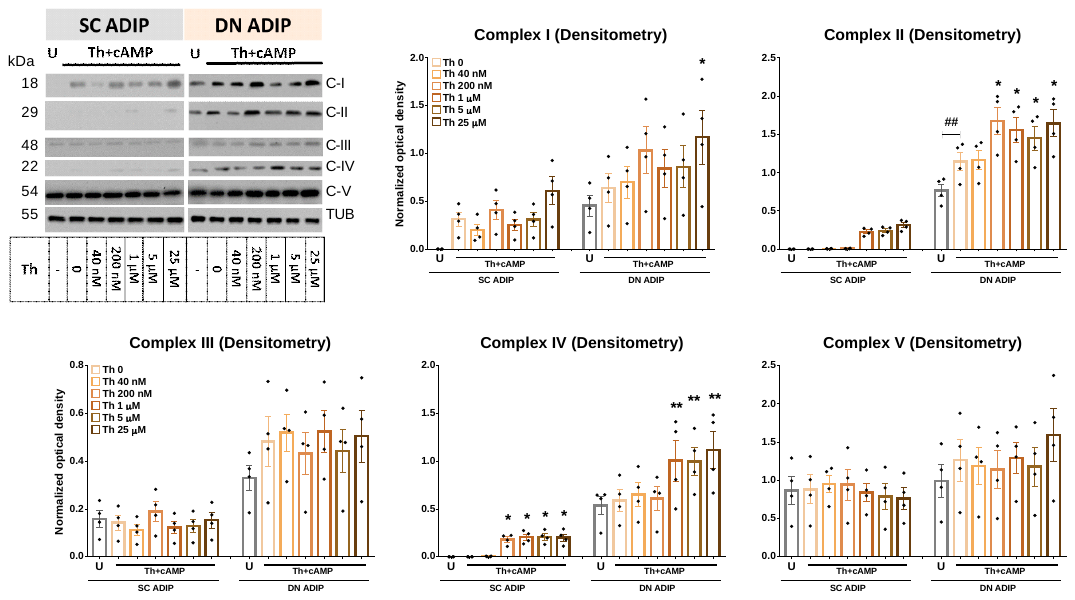

Figure S9. The effect of gradually increasing concentration of thiamine (Th) on the expression of mitochondrial complex subunits. ADIPs were differentiated and treated as in Figure 5. After differentiation, ADIPs were treated with 500 µM dibutyryl-cAMP and gradually increasing concentrations of Th for 10 hours. Quantification of mitochondrial complex I-V subunit protein expression by densitometry normalized to tubulin, n=4. U= untreated. Statistical analysis was performed by paired t-test, *p<0.05, **##p<0.01; * statistical analysis was performed comparing each concentration of Th to the lack of Th or # comparing the indicated groups

**Supplementary Tables**

Supplementary Table 1. Genes primers and probes

| Genes | Assay ID |
| --- | --- |
| SLC19A3 | Hs00228858_m1 |
| SLC19A2 | Hs00949693_m1 |
| SLC25A19 | Hs01001439_m1 |
| UCP1 | Hs00222453_m1 |
| PPARGC1A | Hs01016719_m1 |
| PRDM16 | Hs00537016_m1 |
| CITED1 | Hs00918445_g1 |
| TBX1 | Hs00271949_m1 |
| DIO2 | Hs00255341_m1 |
| GAPDH | Hs99999905_m1 |

Supplementary Table 2. Antibodies used in immunoblotting

| Antibody | Company | Catalog Number | Dilution |
| --- | --- | --- | --- |
| UCP1 | R&D Systems, Minneapolis, MN, USA | MAB6158 | 1:750 |
| SLC19A3 | Novus Biologicals, Centennial, CO, USA | NBP1-69703 | 1:500 |
| SLC19A2 | Abcam, Cambridge, MA, USA | Ab229680 | 1:500 |
| SLC25A19 | Novus Biologicals, Centennial, CO, USA | NBP1-80528 | 1:500 |
| PDHA1 | Invitrogen, USA | 459400 | 1:1000 |
| PGC1α | Novus Biologicals, Centennial, CO, USA | NBP1-04676 | 1:1000 |
| Total OXPHOS | Abcam, Cambridge, MA, USA | ab110411 | 1:1000 |
| TUBULIN | Santa Cruz, USA | sc-5274 | 1:10000 |
| HRP-conjugated goat anti-rabbit IgG | Advansta, San Jose, CA, USA | R-05072-500 | 1:5000 |
| HRP-conjugated goat anti-mouse IgG | Advansta, San Jose, CA, USA | R-05071-500 | 1:5000 |
